# Passage of human-origin influenza A virus in swine tracheal epithelial cells selects for adaptive mutations in the hemagglutinin gene

**DOI:** 10.1101/2025.06.11.659236

**Authors:** Jongsuk Mo, Lucas M. Ferreri, Ginger Geiger, Daniel R. Perez, Daniela S. Rajao

## Abstract

Frequent spillover of influenza A viruses from humans to swine contributes to the increasing diversity of influenza viruses circulating in pigs. Although these events are common, little is known about the adaptation processes that take place when viruses jump between the two species. We examined the changes that occurred during serial passages of a reassortant H3N2 virus (VIC11pTRIG) containing human seasonal surface genes (Hemagglutinin and Neuraminidase) and a swine-adapted internal gene constellation in differentiated primary swine tracheal epithelial cells (pSTECs). The VIC11pTRIG reassortant virus was serially passaged 8 times in pSTECs and compared to a control swine-adapted strain (OH/04p) containing the same internal gene constellation. Viral RNA from passages 0 (inoculum), 1, 3, 4-8 were sequenced via next generation or Sanger sequencing. Hemagglutinin diversity was highest at passage 3. Two amino acid mutations in the Hemagglutinin protein (N165K and N216K) were fixed at passages 7 and 5, respectively. These changes were associated with increased fitness of the virus in pSTECs compared to the original parental strain. Our results suggest that the adaptation of human seasonal H3N2 to swine cells may lead to the selection of HA mutations located near the receptor binding site. These mutations may result in increased fitness of human-origin H3N2 strains to adapt in swine.

## 1. Introduction

Influenza A virus (FLUAV) is a member of the *Orthomyxoviridae* family that causes substantial economic burden to the swine industry. Great genetic diversity is observed between and within the endemic H1N1, H1N2, and H3N2 subtypes that co-circulate in swine globally (1–3). This diversity is largely the result of transmission of FLUAV from other species, followed by antigenic shift and drift in the swine population (4–8). The genetically diverse surface proteins, hemagglutinin (HA) and neuraminidase (NA), of most swine FLUAV circulating in the U.S. currently are paired with different combinations of the triple reassortant internal gene (TRIG) constellation (9), which was introduced to swine in North America several decades ago (10, 11), with genes from the 2009 pandemic H1N1 lineage. FLUAV infects several different animal host species in addition to pigs, aiding in virus dissemination and maintenance. Although FLUAV has the ability to cross between species, sustained transmission in the new host requires adaptive changes to occur (12, 13).

The frequent bidirectional transmission between humans and swine has led to establishment of many novel virus lineages in pigs, contributing to the diversity of FLUAV circulating in this host species (4, 6). Monitoring programs have frequently identified human-origin genes within FLUAV circulating in the U.S. swine population (1, 14, 15), often displaying changes, particularly in the HA, that can alter antigenicity and infectivity in pigs (6, 7, 16). However, the molecular mechanisms that allow human-origin FLUAVs to adapt to pigs are not well understood. Although the HA is recognized as a major determinant of host specificity due to its role in receptor binding (17), other viral proteins have also been shown to affect host specificity (18, 19). Our previous work investigating the adaptation of human-origin H3N2 viruses to pigs showed that although the HA was crucial, the internal gene constellation also played an important role (6), suggesting that the adaptation of FLUAV between humans and pigs is a multigenic trait.

We have previously shown that a reassortant H3N2 virus containing human seasonal HA and NA genes and internal gene constellation from swine-adapted lineages was able to transmit between pigs (20). A critical mutation (A138S) was identified after a single transmission round and shown to improve replication in primary swine tracheal epithelial cells (pSTECs) and in lungs of inoculated pigs (20, 21). These results confirmed that the optimized constellation of internal genes is needed to allow the surface gene segments to persist and evolve in pigs. To test if these results could be recapitulated using an in-vitro system, differentiated pSTECs were used to serially passage the reassortant human-origin H3N2 strain. The pSTECs were grown and allowed to differentiate using an Air liquid interface (ALI) to mimic the natural environment of the host. We identified mutations in the HA gene near the receptor-binding site (RBS) that improved replication in swine cells.

## 2. Materials and Methods

### 2.1 Cells

Human embryonic kidney 293T cells (HEK293T) and Madin-Darby Canine Kidney (MDCK) cells were used for transfection of plasmids for generation of the viruses by reverse genetics. Both cell lines were cultured at 37°C with 5% CO_2_ using Dulbeccòs modified Eaglès medium (DMEM, Sigma-Aldrich, St Louis, MO) supplemented with 10% fetal bovine serum (FBS, Sigma-Aldrich), 1% L-glutamine, and 1% of antibiotic/antimycotics (ab/am; Life Technologies, Waltham, MA). Primary swine tracheal epithelial cells (pSTECs) were used for serial passage experiments and viral growth kinetics. pSTECs were harvested from trachea samples of 5 months-old, influenza-negative female pigs kindly provided by the University of Georgia, College of Agriculture. Briefly, trachea tissues were digested in DMEM/F12 media (Thermo- Fisher Scientific, Waltham, MA) containing 1.5mg/ml of pronase (Sigma-Aldrich, St) and 5% ab/am (Life Technologies) at 4°C for 24 h. Digested cells were washed twice with DMEM/F12 media containing 10% FBS (Sigma-Aldrich) and 10% ab/am solution and subsequently treated with ammonium-chloride-potassium (ACK) lysis buffer (Life Technologies) and DNase (Sigma-Aldrich, St-Louis, MO) followed by collection of pSTECs as previously described (22).

pSTECS were cultured and differentiated under Air-Liquid-Interface (ALI) conditions using collagen-coated (Thomas Scientific, Swedesboro, NJ) transwell inserts (Corning, New York, NY). Cells were cultured in TEC plus media as previously described (22) by adding 500μl of media to the apical well and 1ml to the basolateral portion at 37°C and 5% CO_2_. Trans-epithelial electrical resistance (TEER) was measured to ensure confluency of the cells using the EVOM meter (World Precision Instruments, Sarasota, FL). Cells were cultured in ALI for a minimum of 3 weeks at 37°C with 5% CO_2_. The basal media of the transwell inserts was replaced every 48 h. Cilia activity was checked every 2-3 days .

### 2.2 Viruses

The wild type (wt) A/Victoria/361/2011 H3N2 strain (VIC/11) was kindly provided by Dr. Richard Webby from St. Jude Children’s Research Hospital. The swine-origin wt virus A/turkey/Ohio/313053/2004 (OH/04) was used as the source for TRIG genes and as a control for in vitro studies. The two reassortant viruses, VIC11pTRIG and OH/04p, were described previously (20). The VIC11pTRIG virus carries the HA and NA from VIC/11, the M from A/California/04/2009 H1N1 (CA/09), and the remaining five genes from OH/04. The OH/04p virus encodes the M from CA/09 and the remaining seven genes from OH/04. Three mutant viruses were generated by reverse genetics containing the same internal gene backbone as VIC11pTRIG and OH/04p: VIC11pTRIG_N165K, VIC11pTRIG_N216K, VIC11pTRIG_N165K/N216K (double mutant). The three VIC11pTRIG mutant viruses were designed to carry the same genes as VIC11pTRIG except for the N165K and/or N216K substitutions in the HA (H3 numbering), which were detected during this study.

Mutants were generated by using the Phusion Site-Directed Mutagenesis kit (Thermo- fisher, Waltham, MA,) with specific mutagenesis primers (Table 1). Each primer pair was designed according to manufacturer’s instructions and was used to introduce mutations into the pDP2002_A/Victoria/2011_HA plasmid, which contained the wt HA from A/Victoria/361/2011.

**Table 1.**
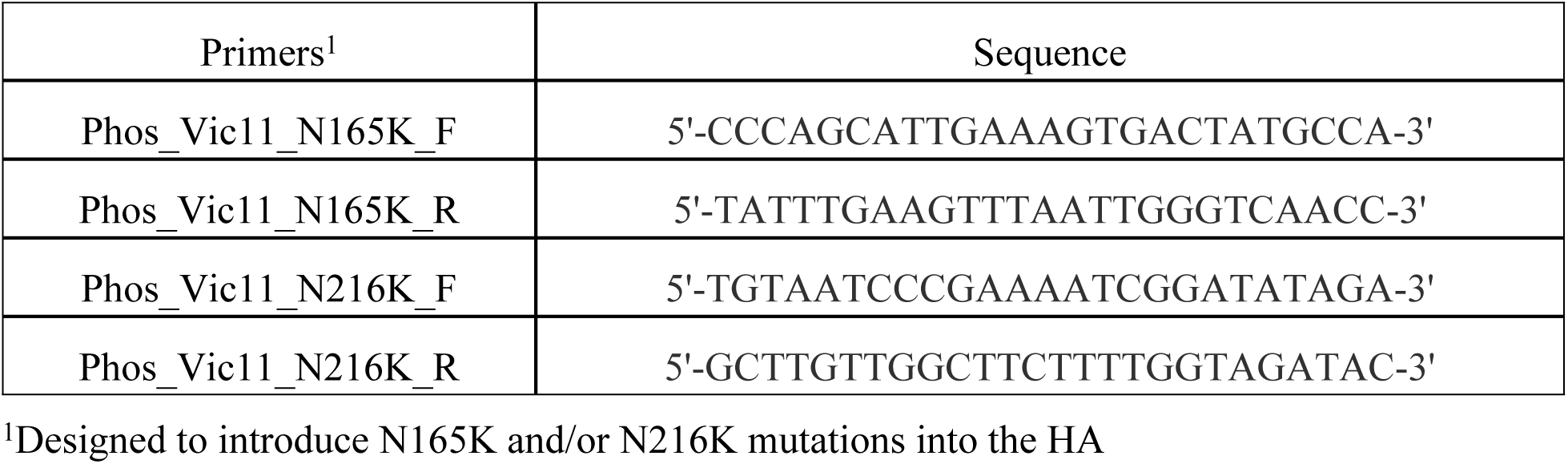
List of mutagenic primers used to generate the VIC11pTRIG variants in this study.

Reverse genetics was performed to generate mutants using an eight-plasmid reverse genetic system and the bidirectional plasmid vector pDP2002 as previously described (23). After transfection, sequences were confirmed by Sanger sequencing (Psomogen Inc, Rockville, MD). Viruses were propagated in MDCK cells for stock preparation.

### 2.3 In-vitro serial passage and growth kinetics

VIC11pTRIG and OH/04p were serially passaged in pSTECs to evaluate the evolution of the HA and NA gene segments. Prior to passage, pSTECs were grown and differentiated for 3 weeks under ALI conditions as described above. For the first passage, VIC11pTRIG was inoculated at a multiplicity of infection (MOI) of 0.1 and OH/04p was inoculated at a MOI of 0.01 in Opti-MEM containing 1% ab/am and 1μg/ml of tosylsulfonyl phenylalanyl chloromethyl ketone (TPCK)-treated trypsin (Worthington Biochemicals, Lakewood, NJ) (infection media). Subsequent passages were conducted by diluting 1/10 of the supernatant from previous passage in infection media. After infection, the plates were incubated for 72 h at 37°C and 5% CO_2_, and CPE was checked daily. After 72 h post inoculation (hpi), viruses were harvested by adding 300μl of infection media to the apical well and gently pipetting to ensure full collection of viruses.

Viral growth kinetics were conducted to evaluate replication of mutant viruses (VIC11pTRIG_N165K, VIC11pTRIG_N216K and VIC11pTRIG_ N165K/N216K) in pSTECs and MDCK cells. For each experiment, VIC11pTRIG and OH/04p were used as controls. Viruses were inoculated into pSTECs or MDCKs with an MOI of 0.05 in infection media for 1 hour at 37°C and were washed once with phosphate buffered saline (PBS) to remove residual virus. Supernatants were collected at timepoints 0, 12, 24, 48, and 72 hpi and samples were stored at -80°C until used for titration via qRT-PCR method as described below.

### 2.4 Virus Titration

Virus titration of serial passage supernatants was conducted using standard TCID_50_. Briefly, 80-90% confluent MDCK cells were inoculated with 10-fold dilutions of the respective viruses in infection media and incubated for 72 h at 37°C and 5% CO_2_. Endpoint titers were read by HA assay with 0.5% turkey red blood cells and calculated by the Reed–Muench method (24).

For titration by qRT-PCR (25, 26), RNA was extracted via the MagMAX™-96 AI/ND Viral RNA Isolation Kit (Thermo-Fisher, Waltham, MA) following manufacturer instructions. For qRT-PCR, the qScript XLT One-Step RT-qPCR ToughMix kit (Quantabio, Beverly, MA) was used according to manufacturer’s instructions, by adding 0.5μl each of forward (5’- AGATGAGTCTTCTAACCGAGGTCG-3’) and reverse primers (5’- TGCAAAGACACTTTCCAGTCTCTG-3’) targeting the FLUAV M segment, 1μl of probe (5’-/56-FAM/TCAGGCCCCCTCAAAGCCGA/36-TAMSp/-3’), 5 μl of RNA, and Nuclease free water up to 15μl per sample. PCR conditions were as follows: 61°C for 30s, 95°C for 30s, then 45 cycles of 95°C for 10s, 60°C for 20s, and 72°C for 10s, followed by final cooling stage of 4°C for 10s using the Quantastudio3 cycler (Thermo-Fisher Scientific, Waltham, MA). Viral titers were calculated based on a TCID_50_ equivalent (TCID_50_eq/ml) using a standard curve of each virus with known titers.

### 2.5 Next-Generation sequencing and Variant Analysis

Virus RNA samples were extracted from tissue culture supernatants either via the MagNA Pure LC RNA Isolation Kit (Roche, Indianapolis, IN) or MagMAX™ AI/ND Viral RNA Isolation Kit (Thermo-Fisher Scientific, Waltham, MA, USA) according to manufacturer’s instructions. Multi-segment, one-step RT-PCR (MS-RT-PCR) was conducted for the amplification of the whole FLUAV genome with the Superscript III High-Fidelity RT-PCR Kit (Thermo-Fisher Scientific, Waltham, MA), as previously described (20, 27). Libraries were prepared using the Nextera XT DNA Library Preparation Kit (Illumina Inc., San Diego, CA), purified, and size-selected using Agencourt AMPure XP Magnetic Beads (Beckman Coulter Life Sciences, Indianapolis, IN) following manufacturer’s recommendation. Collected samples were normalized to 4 nM and pooled after fragment size distribution was analyzed using the High Sensitivity DNA Kit (Agilent, Santa Clara, CA). Pooled libraries were loaded and sequenced using the 300-cycle MiSeq Reagent Kit v2 (Illumina, San Diego, CA). Full genome assembly was done using a previously published pipeline (27). LoFreq 2.1.3.1 (28). was used for variant calling. Based on replicate sequence runs for the same control samples, a frequency threshold of 0.02 was used, with a minimum depth of coverage of 100 and a central base quality score of Q30 or higher. Viral diversity quantification was done using the consensus sequences and single nucleotide variants derived from the variant calling data with the DNASTAR Lasergene® bioinformatics package Version 17 (DNASTAR, Madison, WI).

### 2.6 Visualizing and mapping HA protein structure

The sequences of the HA proteins of the VIC11pTRIG variants identified in this study were converted to .fasta files using DNASTAR Lasergene® Version 17 (DNASTAR, Madison, WI) and ExPASy (Swiss Institute of Bioinformatics resources) (29). The generated protein sequences were submitted to the I-TASSER (University of Michigan, Ann Arbor, MI (30) and visualized with CHIMERA 1.141 (University of California, San Francisco, CA) (31) for their three-dimensional (3D) structures.

### 2.7. Sequence analysis of HA proteins from public databases

The HA sequences from all swine H3 FLUAV from January 2000 to December 2023 were downloaded from the Bacterial and Viral Bioinformatics Resource Center (BV-BRC) (32) and the Global Data Science Initiative (GISAID) (33), to evaluate amino acid residues found at the 165 and 216 positions. HA protein sequences were obtained from 4619 viruses from North America and 1055 viruses from Asia, Europe, South America, and Oceania. HA protein sequences were aligned, and logos were created using WebLogo 3 (https://weblogo.threeplusone.com) (34).

### 2.7. Statistical Analysis

All statistical analyses were conducted using the GraphPad Prism version 9.3.1 (GraphPad, San Diego, CA), with a P-value ≤0.05 considered significant. Statistical methods include nonparametric one-way analysis of variance (ANOVA), two-way ANOVA and nonparametric t-tests. Results shown to be significant were subjected to pair-wise comparisons using the Tukey-Kramer test or the Dunn’s test with Bonferroni correction.

## 3. Results

### 3.1 Serial passage of VIC11pTRIG in pSTECs increased viral replication

Supernatants were collected from passages 1 to 8 (P1 to P8) after serial passages of VIC11pTRIG and OH/04p in pSTECs and titrated by the TCID_50_ method (Fig 1). The viral titers of VIC11pTRIG were relatively low in early passages (P1 and P2) with an average titer of 5.0×10^2^ TCID_50_/ml (Fig 1A). However, titers gradually increased until reaching 1.2×10^5^ TCID_50_/ml in P4, remaining high in subsequent passages, with an average titer of 1.0×10^5^ TCID_50_/ml. In contrast, titers of OH/04p were high from the start of the serial passage experiments, with viral titer of 8.6×10^6^ TCID_50_/ml in P1 and an average of 9.6×10^6^ TCID_50_/ml throughout P8 (Fig 1B).

**Fig 1.**
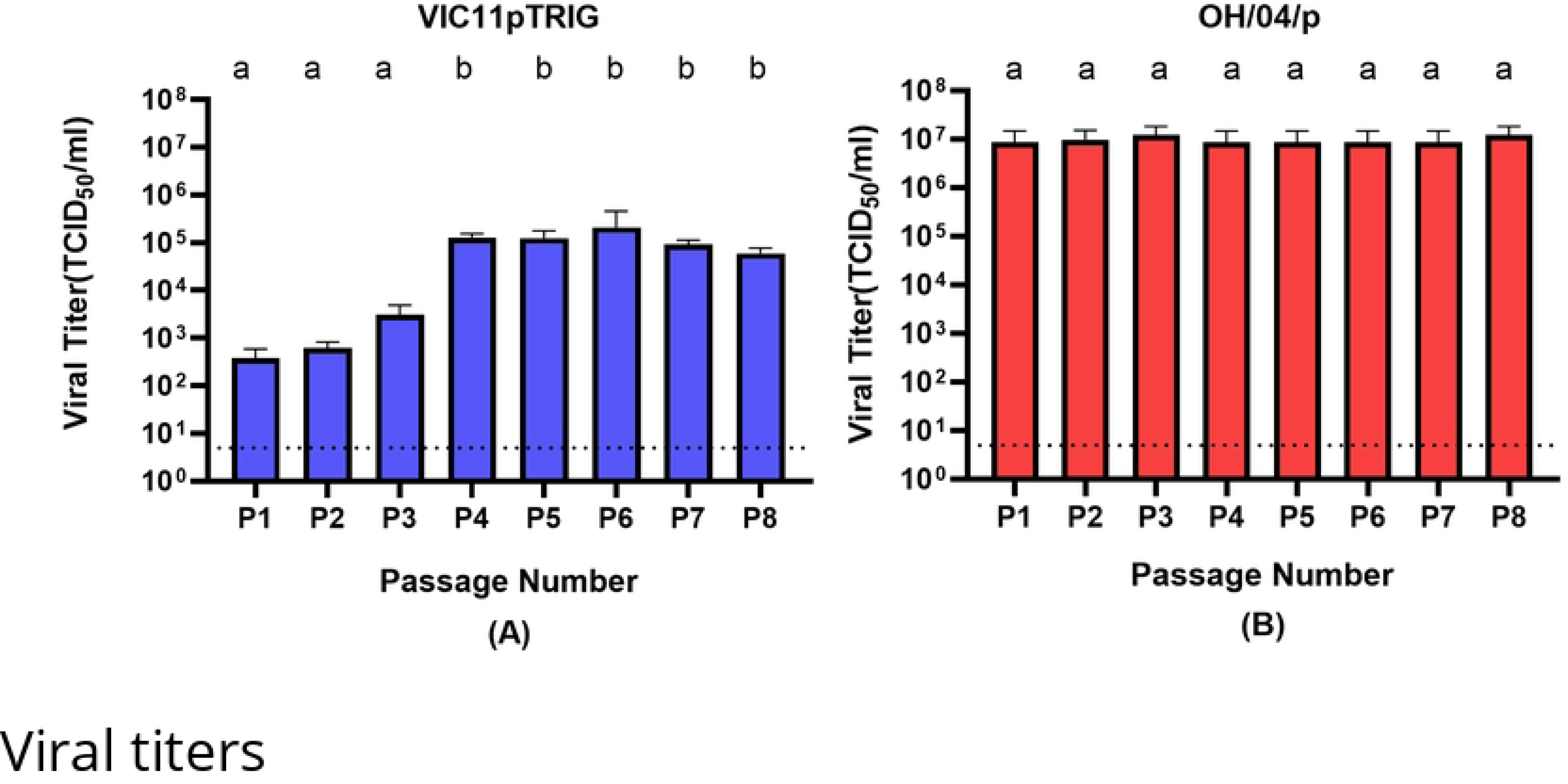
**Viral titers during serial passages of reassortant VIC11pTRIG and OH/04p influenza viruses in primary swine tracheal epithelial cells (pSTECs).** A total of 8 serial passages were conducted in pSTECs. (A) Titers of passage 1 to passage 8 (P1-P8) of VIC11pTRIG virus. (B) Titers of P1-P8 of OH/04p virus. Titers were calculated via the Reed-Muench method. Values are shown as mean TCID_50_/ml titers ± standard error of the mean. Dotted line represents the limit of detection.

### 3.2. Serial passage of VIC11pTRIG in pSTEC resulted in the emergence of dominant variants

To evaluate if changes in viral titers were caused by any changes in viral genotypes, the supernatants from passages 1, 3, 5, and 7 were sequenced by NGS and analyzed for HA and NA nucleotide changes. The supernatants from passages 4, 6, and 8 were sequenced by Sanger sequencing. A total of 12 variants were observed in the HA and 7 variants in the NA of VIC11pTRIG, including a synonymous mutation at nucleotide position 385 (Table 2). Two HA variants, N165K and N216K, became dominant at P4 of VIC11pTRIG (confirmed by consensus Sanger sequencing) and became fixed at P5 (N216K) or P7 (N165K) (H3 numbering) (Table 2). The two dominant mutations were maintained until the last passage (P8). No dominant mutations were detected at P1 and P3 in the HA of VIC11pTRIG. For the OH/04p strain, a total of 14 variants were discovered in the HA and 16 variants in the NA, including two silent mutations emerging at nucleotide positions 240 and 538 at P7 in the HA (Table 2). Only the two silent mutations became dominant for OH/04p (P7).

**Table 2.**
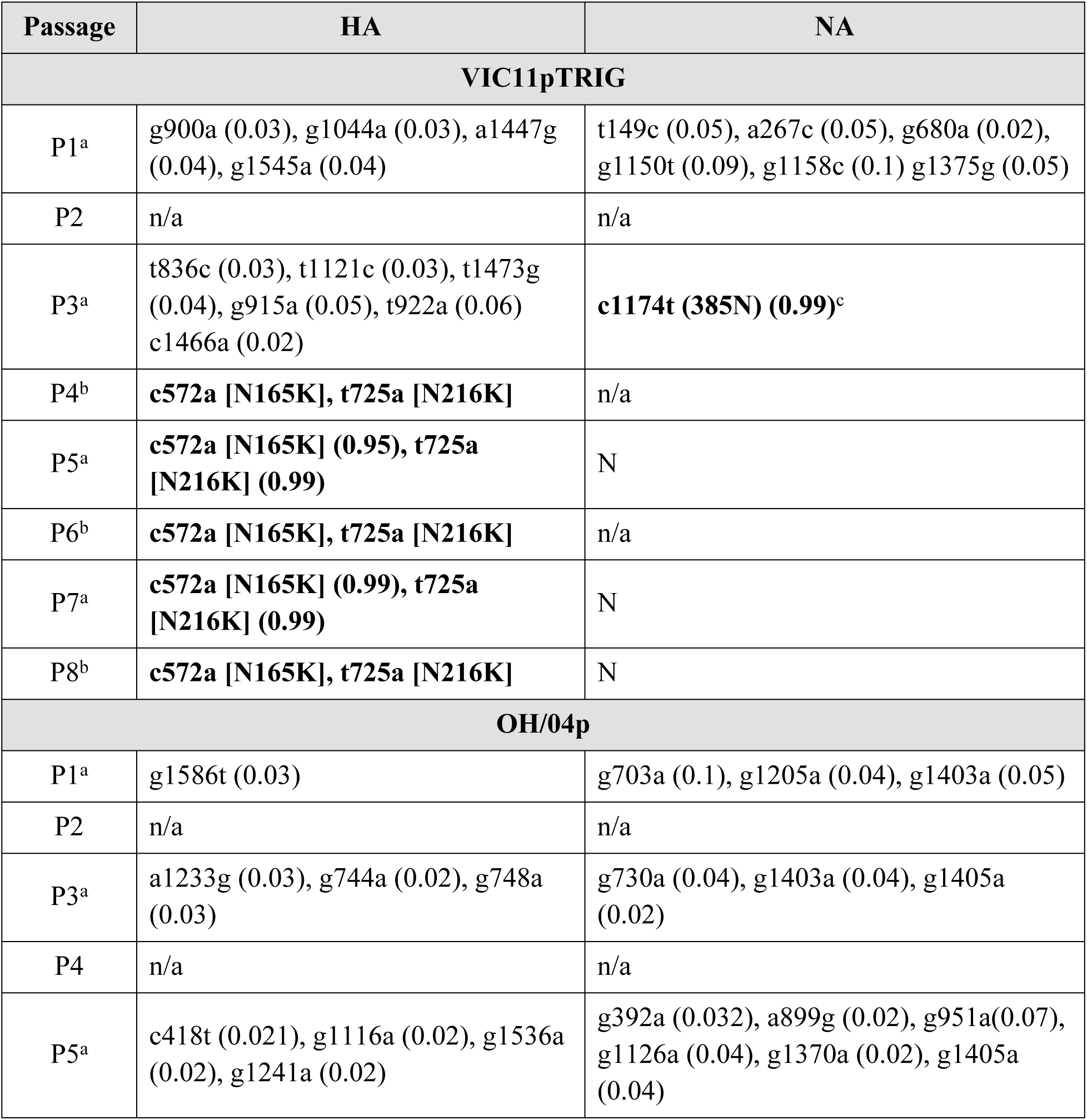
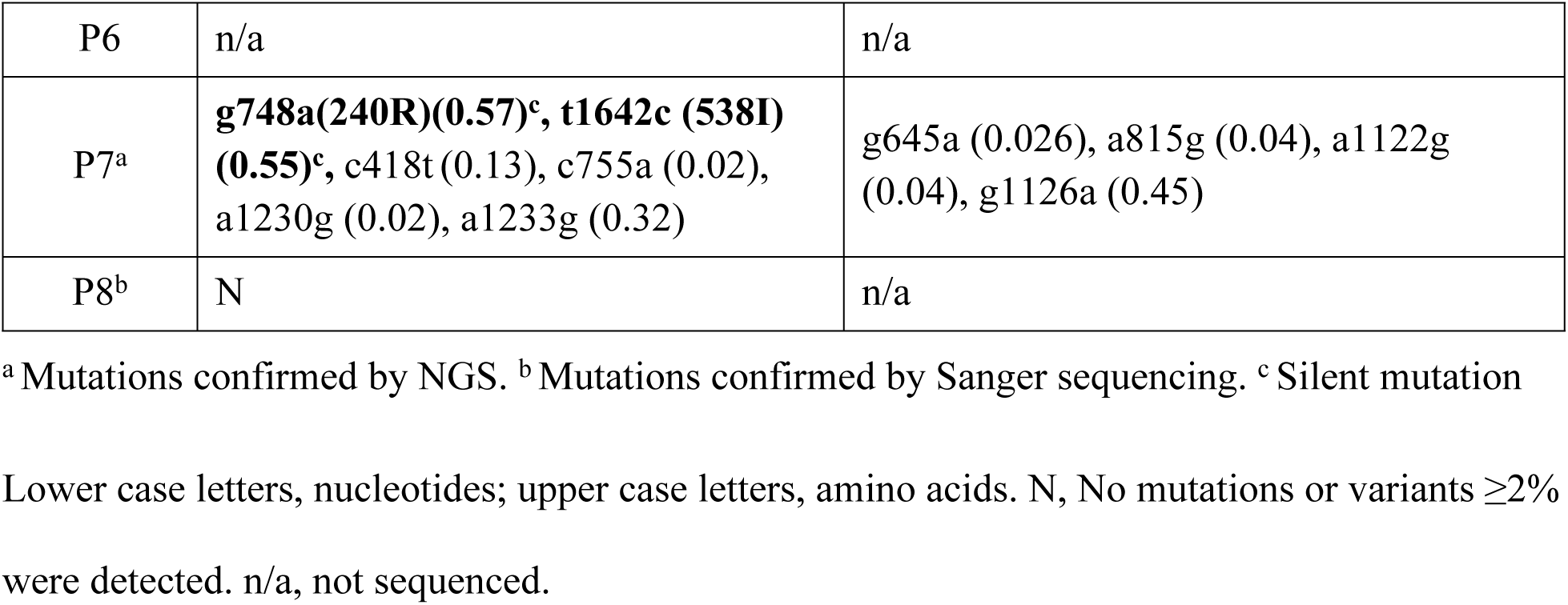
Variants identified during serial passage of VIC11pTRIG and OH/04p in primary swine tracheal epithelial cells (pSTECs). Nucleotide mutations present in ≥2% of variant viruses are shown. All mutations were confirmed by NGS or Sanger sequencing. P1-P8 represent passages 1 to 8. Numbers between parenthesis represent frequency of detection by NGS. Samples sequence by Sanger show consensus changes. Changes in bold represent significant mutations (>50%).

### 3.3. Major variants identified in the HA were located within the receptor binding sites (RBS)

The two dominant variants identified in the HA of VIC11pTRIG were located in the head domain, near the receptor binding site (RBS) (Fig 2). The N165K was located near the 130 loop. The N216K mutation was located between the 130 loop, the 190 helix, and the 220 loop. The two mutations were also located in the H3 antigenic site B (35).

**Fig 2.**
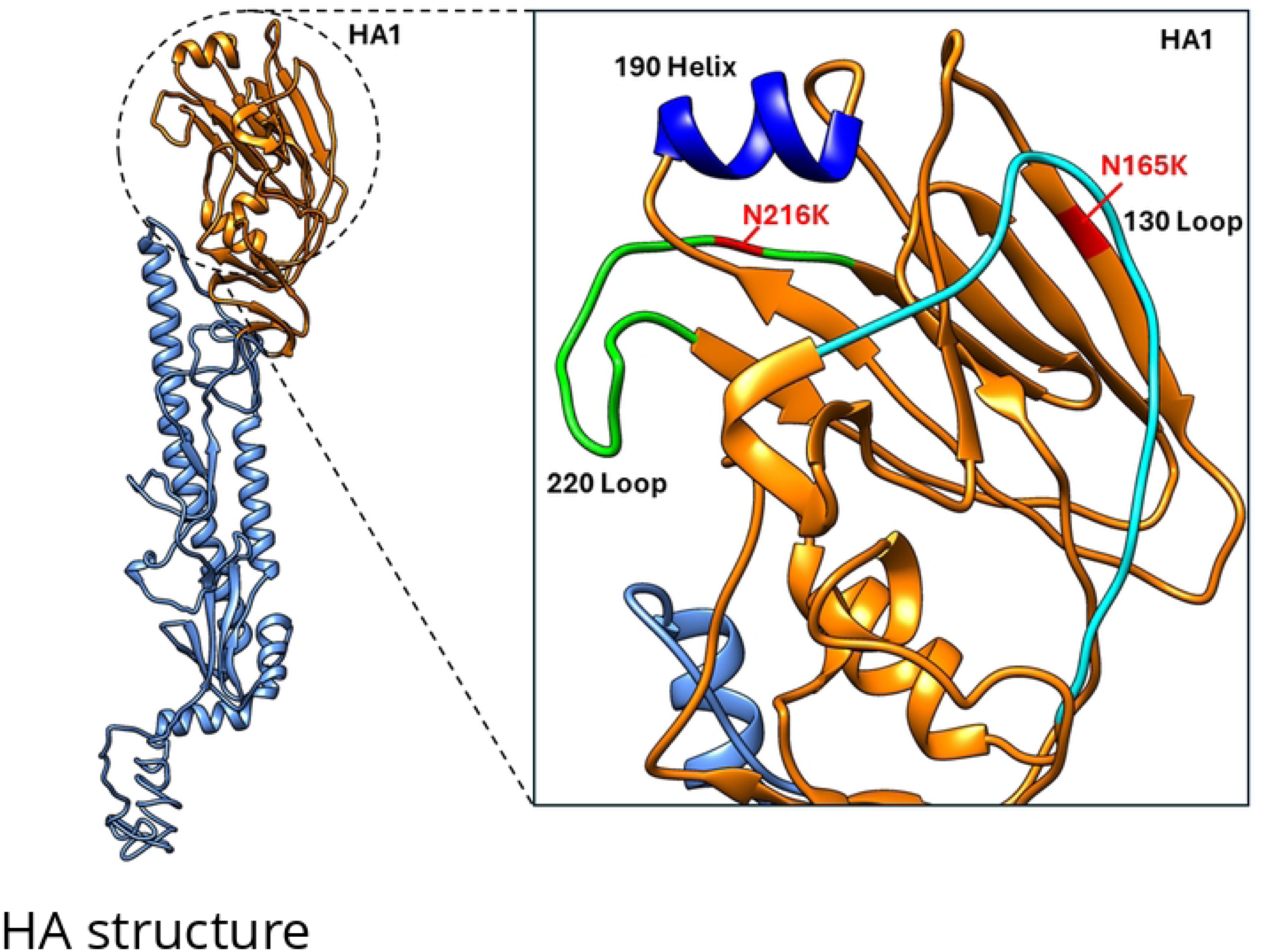
Location of mutations identified on the HA of Vic11pTRIG virus. Dominant mutations identified in this study on the HA of VIC11pTRIG virus are highlighted on the structure of the human A/Victoria/361/2011 hemagglutinin. Red, residues N165K and N216K. The 190 helix, 130 loop, and 220 loop of the receptor binding site (RBS) are shown.

### 3.4 Viruses containing substitutions N165K and/or N216K show better replication than VIG11pTRIG in pSTECs

Mutant strains of VIC11pTRIG were generated containing substitutions N165K and/or N216K and viral growth kinetics were evaluated in pSTECs and MDCKs to compare replication efficiency in relation to the original VIC11pTRIG containing wild type HA (Fig 3). Mutants were generated with single or double mutations. Overall, mutants showed more efficient replication compared to the VIC11pTRIG in pSTECs (Fig 3A). The double mutant (VIC11pTRIG_ N165K/N216K) showed the highest titers at 12 hpi in these cells, with average titer of 1.2×10^5^ TCID_50_eq/ml, followed by VIC11pTRIG_N165K and VIC11pTRIG_N216K, with average titers of 1.1×10^4^ TCID_50_eq/ml. The OH/04p and VIC11pTRIG strains showed the lowest titers at this timepoint. While the VIC11pTRIG strain titer remained relatively constant after 12 hpi, all other viruses continued to grow and remained higher than VIC11pTRIG in the subsequent timepoints. At 48 and 72 hpi, VIC11pTRIG_N216K and OH/04p showed similar titers at an average of 8.5×10^6^ TCID_50_eq/ml titer, followed by VIC11pTRIG_N165K and VIC11pTRIG_ N165K/N216K, at an average of 3.4×10^5^ TCID_50_eq/ml, all significantly higher than VIC11pTRIG by at least 2 logs. In contrast, there were no significant differences observed among the viruses’ growth kinetics in MDCK cells (Fig 3B).

**Fig 3.**
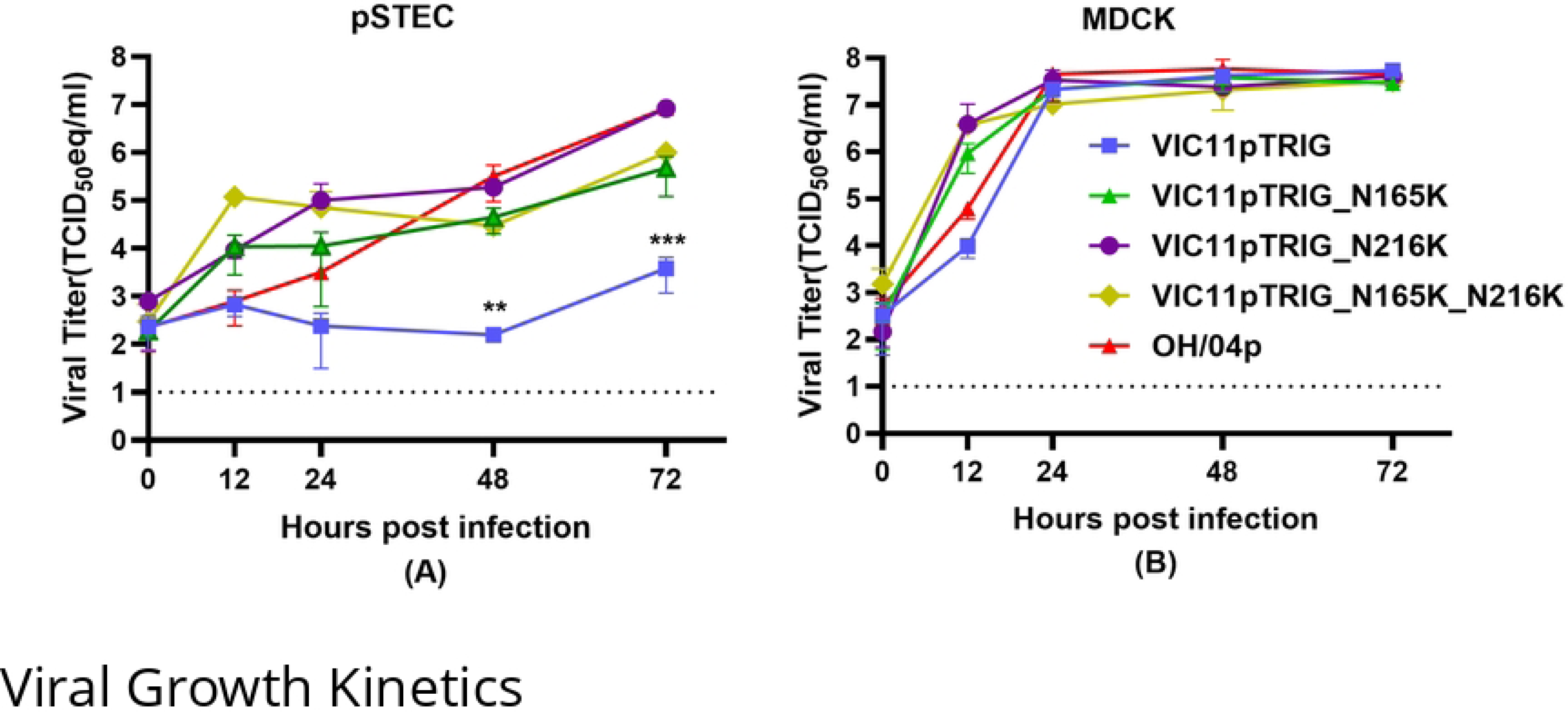
In vitro replication kinetics of mutants containing N165K and/or N216K substitutions in the HA of VIC11pTRIG in primary swine tracheal epithelial cells (pSTECs) and MDCK cells. Viral growth kinetics of mutant viruses (VIC11pTRIG_ N165K, VIC11pTRIG_N216K, VIC11pTRIG_N165K/N216K), VIC11pTRIG, and OH/04p in (A) pSTECs and (B) MDCK cells. pSTECs and MDCK cells were infected at an MOI of 0.05, and supernatants were collected at 0, 12, 24, 48, and 72 h post infection (hpi). Viral titers were quantified by qRT-PCR and titers calculated by TCID_50_ equivalency. Values are shown as mean TCID_50_/ml equivalent titers ± standard error of the mean. Dotted lines indicate limit of detection.

### 3.5. FLUAVs isolated from pigs globally show evolution in the 165 and 216 positions in recent years

To assess the amino acids at the 165 and 216 positions found in natural isolates detected in pigs globally, a total of 5829 HA protein sequences were obtained from the BV-BRC/GISAID. There was minimum variation at position 165 between the years 2000 and 2006 (Fig 4A-B) in viruses isolated globally. However, between the years 2015 and 2022, a significant percentage of viruses containing Glutamine (E) started to be detected in North America, with prevalence increasing to about half of the viruses collected from 2023 and beyond (Fig 4A). For viruses isolated from Asia, Europe, South America, and Oceania after 2007, Lysine (K) was detected in a smaller percentage of strains compared to Asparagine (N) (Fig 4B). There was very minimum variation at the 216 amino acid position from viruses isolated globally, with most strains showing Asparagine (N) at this position. However, strains from other regions outside North America showed some variation after 2014, with either Asparagine (N) or Serine (S) (Fig 4C-D).

**Fig 4.**
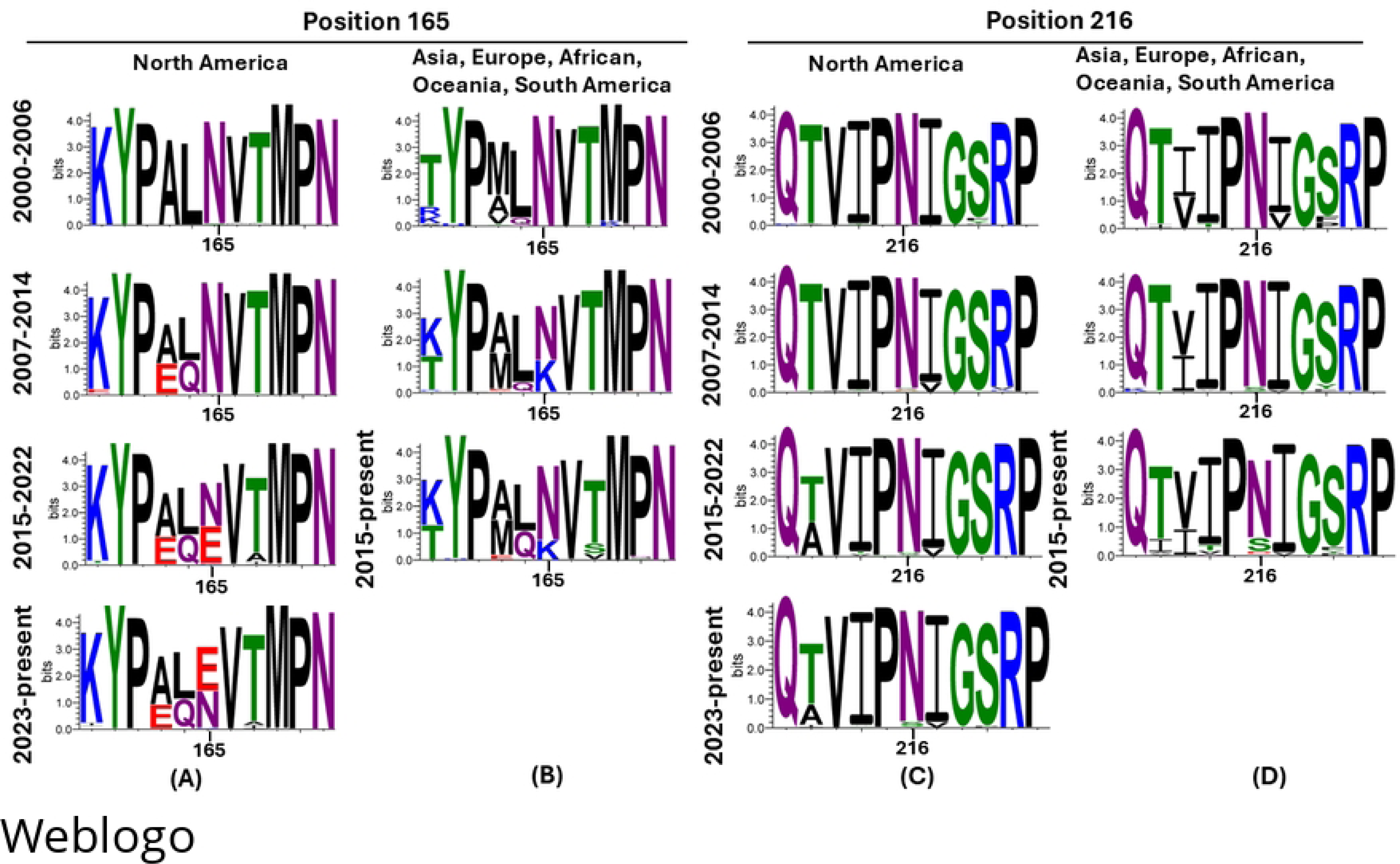
Amino acid substitutions at position 165 and 216 on the HA from H3N2 viruses isolated from pigs globally between years 2000-2025. The relative proportion of mutations between residues 160 and 170 position and 211 and 221 are shown in the figure, staggered for every 7-8 years. Variations in position 165 in swine H3N2 FLUAVs isolated between 2000-2025 in (A) North America and (B) Asia, Europe, Oceania, and South America. Variations in position 216 in swine H3N2 FLUAVs isolated between 2000-2025 in (C) North America and (D) Asia, Europe, Oceania, and South America. Panels generated using WebLogo 3.

## 4. Discussion

Human-to-swine spillover events are common, contributing to the large diversity of FLUAV circulating in pigs globally (5, 36, 37). While the factors that allow human-origin FLUAV to adapt to pigs are not fully understood, the optimal combination of genes seems to be an important step. We have previously shown that the genes from the TRIG lineage in combination with the H1N1pdm09 matrix gene formed a backbone fit for replication and transmission of an H3N2 FLUAV containing surface genes of human-origin in pigs (20). Indeed, the permissiveness of the TRIG constellation to varied HA and NA surface proteins, and its efficient replication in pigs, may have contributed to the establishment of human-origin FLUAV lineages in pigs over the years (5, 6, 38). Here, we assessed the evolution of these previously studied human-origin HA and NA during serial passages in pSTECs. Differentiated cell cultures such as pSTECs retain key characteristics of the pseudo-stratified epithelium (39, 40) and have been widely used as a tool for the replacement of animals for cancer research, studies on pharmacology, respiratory toxicology, and infectious diseases such as influenza (39, 41–44).

Here, differentiated pSTECs were used to simulate long term evolution of human-origin influenza FLUAV in swine and evaluate the viral evolutionary dynamics during adaptation.

Two dominant amino acid substitutions (N165K and N216K) emerged in the HA of the human seasonal reassortant virus (VIC11pTRIG) during serial passages in pSTECs and were maintained up to 8 passages. Early passages of VIC11pTRIG did not show efficient replication in swine cells. However, once mutations N165K and N216K became dominant, a significant improvement in virus replication was apparent. The mutations found here differ from the mutation that was previously observed by us after replication of the VIC11pTRIG in pigs (20), likely due to differences in the systems used. Here we used an *in vitro* system in which differentiated cells were derived from animals at 5 months of age (∼20 weeks old), while the animals used in the *in vivo* study were 4-weeks-old, likely incurring age-related individual variation. Additionally, although differentiated epithelial cells are an advantageous system that closely resembles the animal’s respiratory tract, they may lack intrinsic factors and other cells that may be involved during FLUAV infection. Hence, the human FLUAV would have to overcome additional selective pressures during replication and transmission *in vivo* that may not be present *in vitro*. Nevertheless, all mutations were located near the receptor binding site of the FLUAV HA, which suggests that potential receptor differences between the two hosts may impose selective pressures on the virus populations.

The 165 position is one of the most conserved glycosylation sites in H3 FLUAV sequences (45) and was shown to be involved in immune escape in humans (46). N-glycosylations are known to promote immune escape by physically preventing antibodies from binding to specific antigenic sites but they can also shield the receptor binding site and result in decreased binding affinity to cell receptors (46, 47). The N165K mutation in the A/VIC/11 HA leads to a putative glycosylation loss, which may have resulted in improved binding to swine cells. Interestingly, loss of the glycosylation site at position 165 was shown to increase replication of human H3N2 FLUAV in guinea pigs (48). The number of glycosylations on the HA head has also been shown to affect susceptibility to the antiviral activity of innate collectins such as surfactant protein D (SP-D): more glycosylations tend to make viruses more susceptible (49, 50). Since porcine SP-D has been shown to inhibit FLUAV more efficiently than human SP- D, it is possible that this is an important step during infection of swine cells in which selective pressure is exerted on the human-origin HA.

Little variation existed at position 165 in swine isolates detected globally until 2006, but variation increased between the years 2007 and 2014. For the U.S., this increase in variability coincides with the establishment of the new 2010 H3N2 lineage in pigs (16), an HA lineage that has similar ancestry to the hVIC/11 HA tested here. To some degree, our study recapitulates the evolution observed for position 165 while the novel human-origin 2010 virus was spreading in the swine population. Although the same amino acid substitution was not observed in our study, changes observed in natural isolates in North America also led to loss in glycosylation.

Nevertheless, the K165 amino acid was detected in swine FLUAV strains circulating outside of North America, becoming predominant during a period between 2007 and 2014. Residues in position 216 seemed to be more conserved than in position 165. While the function of the position is not completely known, here the mutation led to enhanced replication in pSTECs to a similar degree of the 165 variant and the double mutant. No additive effects were observed between N165K and N216K substitutions and they showed a similar effect in viral fitness either alone or in combination. Overall, we have identified potential positions that may be involved with adaptation of a virus containing human-origin surface genes to swine. Further studies are needed to evaluate the effect of these mutations in vivo and on subsequent virus evolution.

## Acknowledgments

We thank producers, swine veterinarians, and diagnostic laboratories for providing surveillance data to public databases. We thank the Meat Science Technology Center at the University of Georgia College of Agriculture for providing tissues for cell collection.

## Data Statement

Data will be made available upon request.

